# Illegal hunting threatens the establishment of a lynx population and highlights the need of a centralized judiciary approach

**DOI:** 10.1101/2020.08.16.252890

**Authors:** Raphaël Arlettaz, Guillaume Chapron, Marc Kéry, Elisabeth Klaus, Stéphane Mettaz, Stefanie Roder, Sergio Vignali, Fridolin Zimmermann, Veronika Braunisch

**Author notes:** Corresponding author: Prof. Dr Raphaël Arlettaz, Institute of Ecology and Evolution, Division of Conservation Biology, Baltzerstrasse 6, CH–3012 Bern.

## Abstract

1. Illegal hunting (poaching) represents a major threat to the conservation of large predators. Yet, its impact remains difficult to quantify as there are strong incentives to conceal this criminal activity. Attributing changes in the population status of large carnivores in part to poaching is therefore an important conservation challenge.
2. We present a case study of lynx (*Lynx lynx*) in southwestern Switzerland (canton of Valais) where the current distribution range is much smaller than it was in the recent past and population density is now >80% lower than in other lynx populations in the Swiss Alps, particularly in the adjacent Pre-Alps. We tested four hypotheses to explain this far lower density: 1) a too low density of trail camera-traps deployed for lynx surveys in Valais compared to the Pre-Alps (i.e. a methodological artefact); 2) less favourable environmental conditions around the camera-trap sites; 3) lower densities of the main prey; and 4) poaching.
3. We estimated lynx and ungulate densities and environmental conditions at trail camera sites, and were able to clearly reject the first three hypotheses: 1) the monitoring protocol was similarly effective; 2) environmental conditions around the trapping sites in Valais were even more favourable to lynx detection than in the Pre-Alps; and 3) prey supply was even larger. Concerning hypothesis 4, we discovered a local, but dense network of 17 illegal lynx traps in the narrow main immigration corridor into Valais from the thriving adjacent lynx population in the Pre-Alps, suggesting intense local poaching.
4. Our findings substantiate the suspicions of long-lasting lynx poaching as a threat to the establishment and survival of the Valais population. The fact that instances of poaching were publicly known since 1995 but remained unabated for at least two decades, until a first conviction occurred, questions the commitment of local authorities to address this case of wildlife crime. Our study shows that inquiries about wildlife crime such as top predator poaching may need to be carried out at the highest levels of jurisdiction to avoid any risk of collusion between law enforcement agents and poachers.

## Introduction

The protection of large predators in human dominated landscapes represents an important challenge for contemporary conservation. In particular, the predatory activity of large carnivores may interfere with economic activities, notably domestic animal husbandry, and leisure or commercial hunting ^1^, rendering a smooth coexistence with all humans difficult to achieve ^2, 3^. Although globally large carnivores are declining ^4^, some regions of the world have seen these species making an often unforeseen comeback ^5^. One essential driver of such recoveries, besides the increase of the prey species and a more favourable public opinion, has been the legal protection conferred to these species ^5^. In many countries, it is a crime today to kill large carnivores outside strictly defined conditions approved by the relevant authorities. For example, in the European Union, the Habitats Directive imposes strict limitations to the legal killing of large carnivores ^6^.

However, illegal activities remain a constant threat and recent research has demonstrated that the illegal killing of large carnivores may be much more frequent and widespread than generally thought. In Scandinavia, Liberg et al. ^7^ found that half the mortality of wolves (*Canis lupus*) was due to poaching and that two thirds of this poaching mortality was not directly observable. Andrén et al. ^8^ found that poaching accounted for 46% of the mortality of adult lynx in Norway and Sweden. In Croatia, Sindičić et al. ^9^ investigated mortality causes of lynx and found that poaching accounted for 60% of all recorded fatalities from 1999 to 2013. Heurich et al. ^10^ found that the source-sink dynamics of reintroduced lynx populations in Central Europe was essentially driven by poaching. A survey among Czech hunters revealed that more than one third of them knew of instances of lynx poaching, with 10% even anonymously claiming to be the actual perpetrators ^11^.

Because poaching can be such a substantial source of mortality in lynx populations, documenting its exact magnitude is essential for conservation planning. However, since poaching is illegal by nature, there are strong incentives to hide it from law enforcement authorities, making it very difficult to document and quantify. In addition, sometimes it remains unclear whether law enforcement forces are passively complicit of poaching by failing to prosecute poachers when evidence is available or even by advising poachers on how to evade legal prosecution ^12^.

In this paper, we report long suspected but until recently unpunished poaching activities in what is presumably *the* key lynx dispersal corridor in the southwestern, inner Swiss Alps. This provides evidence that illegal hunting threatens that population which was reintroduced in the early 1980s and was thriving during the initial years. If suspicions of poaching already existed in that region of Switzerland since the time of its reintroduction ^13^, first public evidence of lynx poaching emerged in 1995, when a photograph of a hunter portrayed with a rifle and two dead lynx was broadcast by the media ^14, 15^ (Fig. S1).

However, a local court then accepted the hunter’s claims that he had merely found the animals that were already dead, and acquitted him. More substantial evidence for poaching in this area emerged in 2011 with the discovery by ourselves of a first active lynx trap. This occurred at the start of our long-term research programme aimed at developing simple and cost-effective methods for investigating the spatio-temporal relationships between large carnivores and their ungulate prey, especially in the context of the natural return of the wolf into the Swiss Alps. In this programme, we deployed a network of trail camera-traps to assess the spatial distribution and population densities of large carnivores. This monitoring effort led us to discover unexpectedly low densities of lynx in the southwestern Swiss Alps (Valais), reaching less than 20% of the densities observed in other areas of Switzerland where environmental conditions are fairly similar and where standard lynx monitoring schemes have been carried out for many years ^16-19^. In addition, our programme revealed an almost complete absence of reproduction in Valais ^20^. This low density compromised the research project from the onset (providing too small sample sizes to allow drawing inferences on predator-prey interactions), but it also raised questions about the possible factors causing it.

In this paper, we evaluate four plausible hypotheses that could potentially explain the extremely low density of the lynx population in Valais of only 12-20% of what might reasonably be expected, as framed in a previous assessment of its status ^20^: methodological limitations and caveats linked to the lynx monitoring techniques themselves, suboptimal ecological conditions for the lynx in Valais, and high poaching pressure ^20^. Elucidating the causes of the very low lynx density in Valais is of major conservation relevance for the wider distribution of this felid in the western Alps because the Valais lynx population could act as a key stepping-stone for further recolonisation of the Italian and French Alps where the species is still rare ^13, 20, 21^.

## Material and methods

### Hypotheses

The four hypotheses that are most likely explaining restricted distributional range and low lynx density in Valais were already described in an earlier paper ^20^.

- The first hypothesis proposes that a less dense trail camera-trap network in the Valais project compared to other areas of Switzerland where lynx monitoring is routinely carried out might lead to underestimating local lynx population densities in Valais.
- The second hypothesis states that there are fewer obligate passages for travelling lynx in the landscape configuration of Valais compared to the Pre-Alps. This might induce a reduced probability of lynx photo-capture at the camera-trap sites in Valais.
- The third hypothesis says that lower local densities of the main prey species (roe deer *Capreolus capreolus* and chamois *Rupicapra rupicapra*) in Valais preclude lynx densities from reaching values as high as in other, healthy populations in Switzerland.
- The fourth hypothesis relates to a possibly higher poaching intensity in Valais compared to adjacent regions in Switzerland where lynx thrive. This latter hypothesis, discussed earlier on ^20, 22^, is a legitimate hypothesis to investigate because there is some evidence to suggest that poaching may be not uncommon. This includes a hunter posing with two dead lynx (Fig. S1) and a senator representing Valais in the Swiss parliament officially and merrily claiming in 2010 that the Valais lynx population “*can be managed and regulated in total transparency*” ^23^ – although the lynx is a strictly protected species in Switzerland and no permits for lynx culling or regulation had ever been issued to the Valais authorities by the relevant Swiss federal governmental agency.

### Study area

We conducted this study between 2011 and 2016 in two regions of the Swiss Alps with widely different lynx densities: the Valais Alps (hereafter Valais) in the southwestern Swiss Alps, and the northwestern Swiss Alps (hereafter Pre-Alps) situated on the northern outer margin of the Alpine range, immediately adjacent to the north of Valais ^24^ (Figs. 1 & 12). The Pre-Alps study area comprises parts of the cantons of Bern and Vaud, including Simmental, Diemtigtal, Saanenland and the Pays d’Enhaut, whereas the Valais region is separated by the deep Rhône river valley into the Bernese Alps in the North and the Pennine Alps in the South.

**Figure 1.**
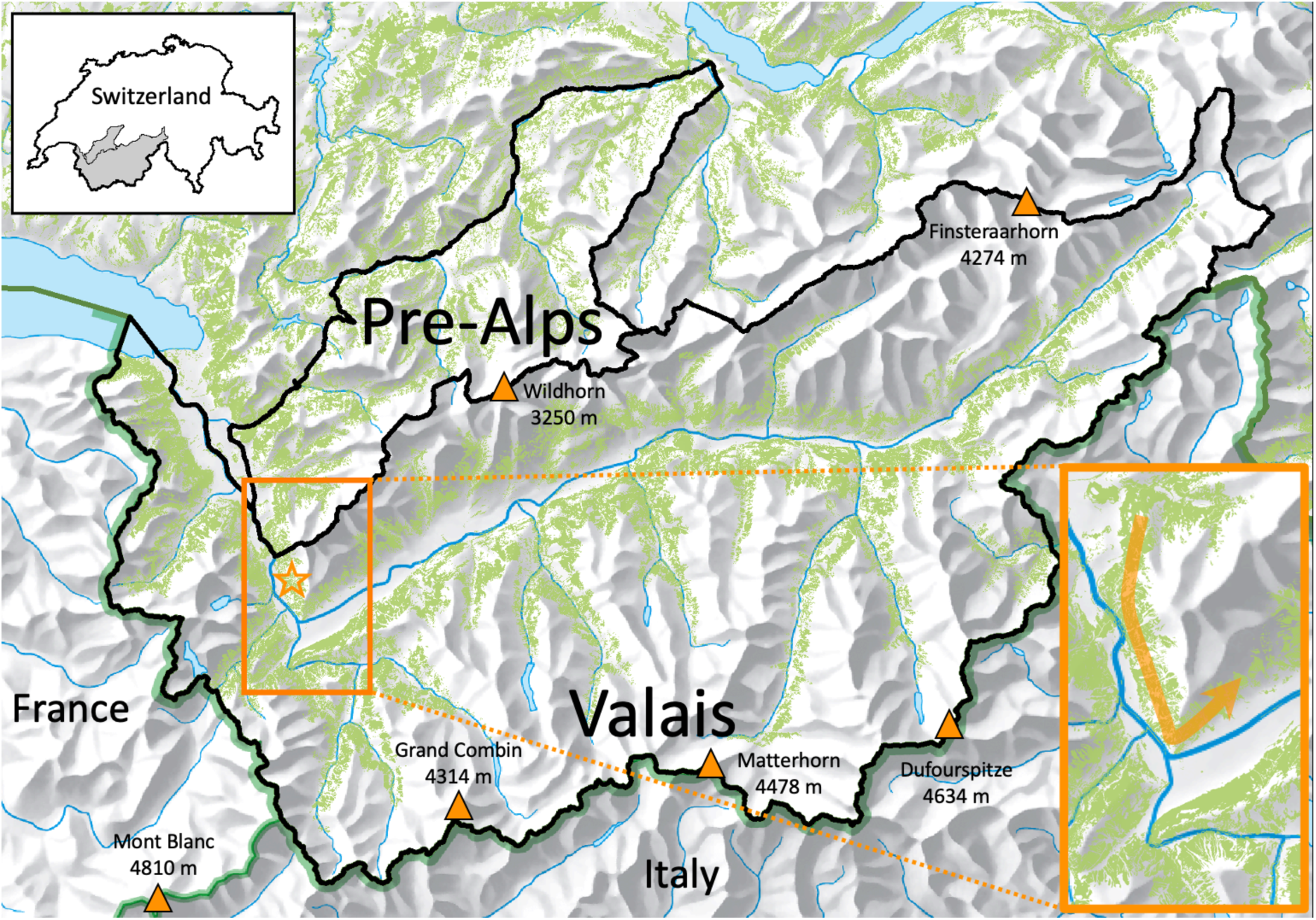
The forest cover continuum in SW Switzerland (green; not shown for France and Italy) depicting the distribution of lynx main habitat. Valais is enclaved within high mountain ranges culminating at more than 4000 m a.s.l., both in the N and S. This isolates the Valais lynx population, with little connectivity to the N (where dense lynx populations occur), E and S (Italy). The forest continuum is higher in W Valais, with an important dispersal corridor linking to the Pre-Alps (Vaud), where lynx thrives, and France, S of the Rhône, where lynx is rare. The main dispersal corridor (orange arrow) into the Upper Rhône valley (Valais) from the thriving lynx population in the Prealps is shown in the inset. The star indicates the location of the network of 17 illegal traps found in the present study (see also Fig. 6).

Although immediate neighbours, the two study areas differ in climate, topography, elevation, and forest cover. The Valais is characterized by a vast altitudinal gradient, from 372 m to 4634 m a.s.l., with the high elevation areas receiving much precipitation, whereas the valley bottoms are particularly arid (ca 550 mm annual precipitation) due to the surrounding high peaks. With a total area of 5224 km^2^, 22.2% are covered by forests, the major lynx habitat, distributed in a belt-like shape along the slopes of the main valley and its numerous tributaries, with a timberline at 1800-2300 m a.s.l. ^20^. Due to the effects of topography, there are more or less continuous forested areas within Valais itself, but without direct connection to the forested areas situated in the North (where denser lynx populations occur), East or South (Fig. 1). The only forested connection with areas outside Valais is in the Northwest (canton of Vaud, with a thriving lynx population ^25^) – constituting the main dispersal corridor for lynx locally – and in the Southwest, in the Haute-Savoie in France, which has only scarce lynx presence. Six species of wild ungulates are present in the area. Red deer (*Cervus elaphus*), roe deer and chamois are abundant and distributed in the whole area, whereas populations of ibex (*Capra ibex*), wild boars (*Sus scrofa*) and non-native mouflons (*Ovis orientalis musimon*) occur more locally.

In the Pre-Alps, the topo-climatic conditions are less extreme, with a larger amount of precipitation more evenly distributed across the altitudinal gradient. This second study area encompasses 1080 km^2^, covering an altitudinal range of 450-2900 m a.s.l., and is covered by forests that are mostly interspersed with grasslands, with a timberline at 1700-1800 m a.s.l. ^26^. The ungulate community is similar to that in Valais, with the exception of the mouflon that is absent. For a typical forest-dwelling species such as lynx, the Pre-Alps thus offer a continuum of suitable (i.e. forested) habitat (Fig. 1).

Hence, in Valais there is a comparatively less isotropic forest matrix, simply because the Upper Rhône valley is surrounded by the highest peaks of western Europe (Fig. 1). This bears clear implications for lynx dispersal: while the Pre-Alps offer only limited barriers to dispersal, the topography of Valais limits any dispersal opportunities into the Upper Rhône valley, and emigration from it. Except from the aforementioned Northwest connection to the Pre-Alps, dispersal would only be possible through high-alpine passes above the timberline, but these are rarely crossed by lynx ^14^.

In both areas we restricted our surveys to an elevational range from the lowest foothills at 500 m up to 2000 m a.s.l., i.e. excluding the valley bottoms with dense human populations and the mountain peaks devoid of forests. This resulted in survey areas of 2555 km^2^ and 957 km^2^ Valais and in the Pre-Alps, respectively.

### Species data

#### Lynx data

Lynx data were obtained from two different, ongoing monitoring schemes and research programmes. Whereas lynx presence and density in the Pre-Alps (and only recently in Valais ^27^) were assessed every second to fourth year by the Carnivore Ecology and Wildlife Management Foundation (KORA) ^28^, lynx density estimates for Valais were obtained from the trail camera network deployed by the Conservation Biology Division of the University of Bern since 2011/12 as part of a study of the spatio-temporal relationships between predators and their ungulate prey.

In the Pre-Alps, 79 trail camera-trapping sites were selected by KORA in a stratified manner in winter, placing a trapping site in every second grid cell of a square raster with 2.7 km side length, resulting in a trap density of 4.7/100 km^2 25^ (Fig. 2). At each location two camera-traps equipped with white flash (Cuddeback, Inc. Green Bay, Wisconsin, USA) were installed (not symmetrically facing each other because of flash interference) in order to facilitate lynx individual identification by assessing the fur pattern on both body flanks ^25^.

**Figure 2.**
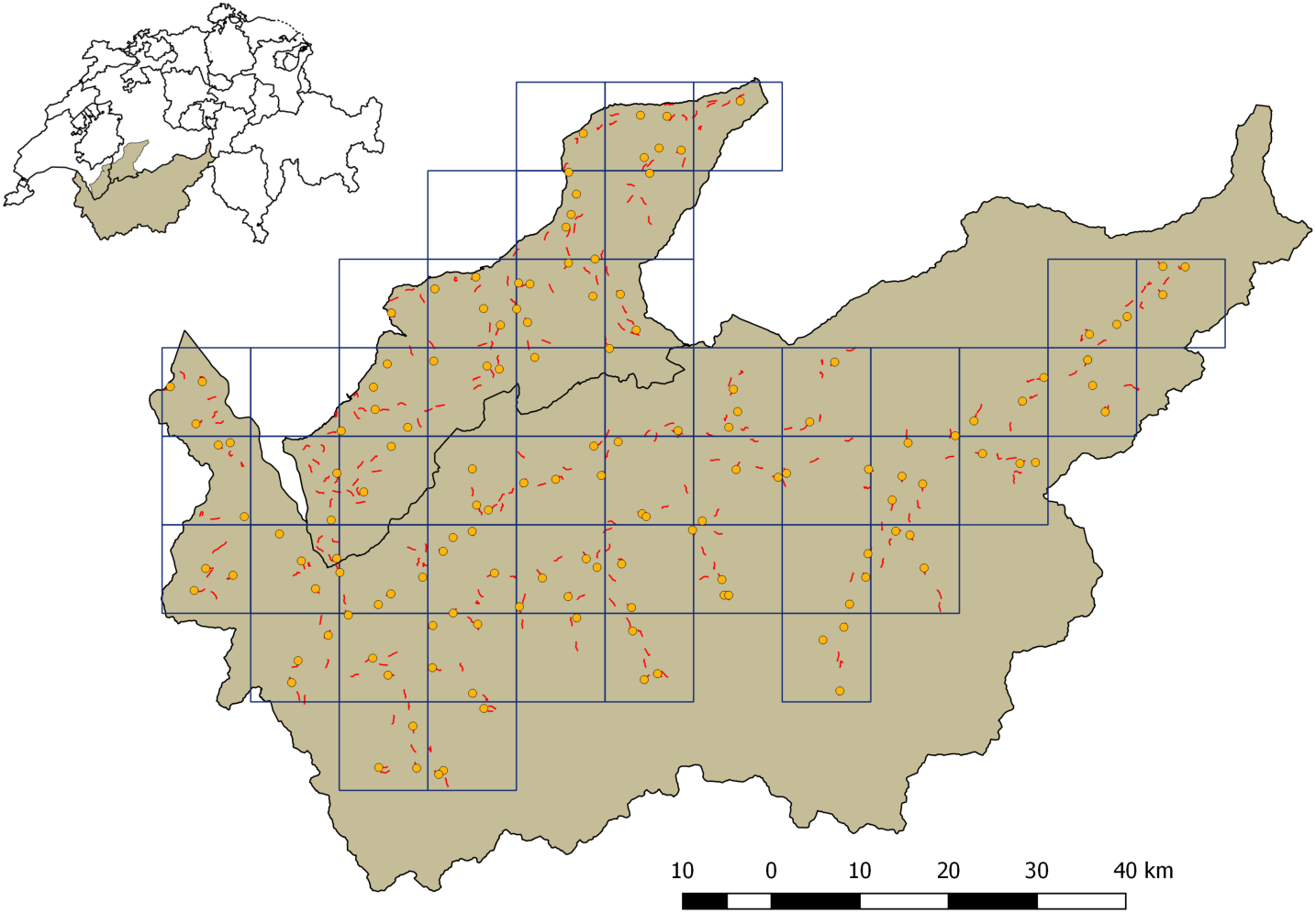
The study areas in the Pre-Alps (upper perimeter) and Valais (lower perimeter) and their location within Switzerland (upper left insert). The grid (10 x 10 km) indicates the raster used for sampling stratification, red lines show the 313 1-km long transects along which snow-tracking was performed. The orange dots depict the locations of the camera-traps.

In Valais, the forested area below 2000 m altitude, corresponding roughly to two thirds (∼3700 km^2^) of the cantonal territory, was divided into 34 grid cells of 100 km^2^ (10 km side-length), each cell being attributed to one of seven topographically defined landscape compartments (Fig. 2; see also Fig. 2 in ^20^). In each of these 34 cells, three camera-traps fitted with a movement and heat detector, and an infrared flash (Reconyx PC900 Hyperfire Professional, Inc. Reconyx, Wisconsin, USA) were placed, resulting in a total of 102 camera-traps, i.e. a camera-trap density of 3/100 km^2 20^. Yet, contrary to the KORA approach, cameras were used singly, not allowing picturing the two flanks of lynx body.

To allow a direct comparison of the camera survey outcomes, the same monitoring scheme as in the Pre-Alps was applied during the winter 2013/2014 to the northwestern part of the Valais study region by KORA, parallel to the ongoing survey scheme by the University of Bern. The comparison of the two simultaneous surveys carried out by the two research groups in the same area delivered information about the influence of trail camera-trap density on the number of individual lynx identified, and therefore lynx density estimates (see ^20^).

In both study areas, camera-traps were placed along forest roads, trails or game paths. Obligate passages, i.e. sites where the movements of animals are strongly constrained by the topography or other natural obstacles, such as large rocks, where chosen wherever possible ^20^. The camera-traps were operational either between mid-November and late March (University of Bern) or January-March (KORA).

#### Ungulate data

Ungulate data were collected using snow tracking along 1-km transects (Valais: n = 218, Pre-Alps: n = 95) in winter 2015/16. In order to place transects in a stratified manner, the two study areas were divided into grid squares with a side-length of 10 km (Fig. 2) (34 and 16 squares in Valais and in the Pre-Alps, respectively) in a way similar to the trail camera sampling design. In each square, on average six transects (range: 1-10) were placed so as to represent the altitudinal and environmental gradients, while accounting for accessibility, topography and human safety with respect to avalanches. Transects were visited twice during each winter (December to March). All tracks found in the snow of the two main prey species of the lynx (roe deer and chamois) were recorded. Based on imprint size and track configuration, we estimated the minimum number of individuals present at each visit. Multiple individuals were counted if tracks of different sizes (i.e. from different sexes or age classes) or individuals travelling together in a group could be distinguished, using a conservative approach: when in doubt, we always recorded the lower number of individuals (for details see ^29^).

### Environmental variables

#### Topographic conditions at the camera-trap sites

To determine the factors influencing detection probability at the camera-trap sites, we recorded 14 variables. Eleven variables were mapped in the field, at the local scale, around the camera sites, while the remaining three were assessed at landscape scale from extant geographical data (Table 1).

**Table 1.** Variables describing the surroundings of the camera-trap sites, obtained from field mapping (local scale) or extant databases (landscape scale). Local scale variables were collected either within a 25-m diameter circular plot around the camera-trap location or along a 25-m long and 2-m broad straight line in the axis of the camera coverage (Fig. S2). See Material and methods for more details.

| Variable | Scale | Description |
| --- | --- | --- |
| **Circular plot** (r = 12.5 m) |  |  |
| Ground vegetation cover | Local | Percentage of ground vegetation cover < 0.5 m |
| Shrub vegetation cover | Local | Percentage of shrub layer cover 0.5–1.5 m |
| Rock cover | Local | Percentage of rock cover |
| Bare ground cover | Local | Percentage of bare ground cover |
| Dead wood cover | Local | Percentage of dead wood cover |
| **Transect** (length = 25 m, width = 2 m) |  |  |
| Slope | Local | In degrees, averaged over transect |
| Barrier | Local | Percentage of total barrier |
| Undergrowth barrier | Local | Percentage of total barrier consisting of undergrowth cover |
| Rock barrier | Local | Percentage of total barrier consisting of rock cover |
| Tree trunk barrier | Local | Percentage of total barrier consisting of tree trunk cover |
| Dead wood barrier | Local | Percentage of total barrier consisting of dead wood cover |
| **Circular window** (r=564 m) |  |  |
| Slope | Landscape | In degrees (continuous) |
| Forest cover | Landscape | Percentage of forest cover (continuous) |
| Road density | Landscape | Percentage of road cover (continuous) |

Topographic and vegetation conditions at all camera trapping sites were assessed between June and August 2016. Variables were acquired within a circular plot (12.5 m radius) directly around the camera location or from a 25 m (i.e. matching that plot diameter) long transect (2 m width) perpendicular to the expected lynx trajectory (along a road or path), i.e. exactly in the axis of the camera coverage (Fig. S2). Within the plot we estimated the vegetation cover of both ground (<0.5 m) and shrub (0.5–1.5 m) layers, rocks, bare ground and deadwood. Along the transect, we estimated the total percentage of barriers, defined as any obstacle that would constrain lynx movement at a height of 0.3– 1 m above ground. In addition, we separately recorded the percentage of dense undergrowth, rocks, dead wood and tree trunks as well as streams wider than 1 m. Slope was measured using a smartphone app (Slope Angle, © Radislav 2015). At a landscape scale, average slope, percentage of forest cover and forest road density were assessed within a radius of 564 m around the camera trapping sites, which corresponds to an area of 1 km^2^. Spatial analyses were performed using QGIS (version 2.18.9) ^30^.

#### Environmental predictors for modelling ungulate densities

The environmental variables we used for predicting area-wide relative ungulate densities (Table 2) were extracted from existing digital information, so as to form basic grid layers with a 25 x 25 m resolution. We obtained data on topography (altitude, roughness, slope, topographic position index) from the Swiss national digital elevation model (DEM). Land cover data (forest, rock, scree, waterbodies, anthropogenic areas, roads and railways, ski lifts) were derived from vector25 ^29^, a digital vector-map regularly updated by the Federal Office for Topography Swisstopo (https://www.swisstopo.admin.ch), except for grassland cover which was obtained from the Arealstatistics of Switzerland (https://www.bfs.admin.ch). Forest type information originating from Landsat5 data was provided by the Swiss Federal Office of Statistics (https://www.bfs.admin.ch). Winter ambient temperature and precipitation came from the Worldclim dataset (http://www.worldclim.org), which was downscaled from a 1-km^2^ raster to a resolution of 100 x 100 m based on the SRTM-V4 digital elevation model following the method described in ^31^. Finally, information about national wildlife reserves (where hunting is legally banned) was obtained from the Federal Office for the Environment (FOEN). All variables were assessed within a 200 m buffer around the transect (corresponding to an area of 50 ha, which approximates the average home range size in winter for the two main ungulate prey species ^32, 33^) by calculating the proportion (boolean variables) or the average (continuous variables) within this area. Variable preparation was performed using QGIS (version 2.18.9) ^30^ and ArcGIS Release 10 ^29^.

**Table 2.** Environmental variables used to predict relative ungulate densities, extracted from buffer zones around the transects or circular moving windows of 50 ha (see Material and Methods for more details). Continuous variables were averaged, while for boolean variables the percentage of cover was calculated.

| Variable | Source | Description |
| --- | --- | --- |
| **Topography** |  |  |
| Altitude | DEM<sup>a</sup> |  |
| Roughness | DEM | Standard deviation of altitude |
| Slope | DEM |  |
| Topographic position index (TPI) | DEM | Concavity vs convexity of site according to <sup>55</sup> |
| **Land cover and land use** |  |  |
| Proportion of forest | Vector25<sup>b</sup> |  |
| Proportion of and distance to anthropogenic areas | Vector25 |  |
| Proportion of rock and screes | Vector25 |  |
| Proportion of grasslands | Arealstatistics | Pastures, meadows |
| Length of main and small roads | Vector25 | Main roads = categories 1-3, small roads = categories 4-6 |
| Distance to roads/rails | Vector25 |  |
| Forest type (coniferous, deciduous, mixed) | BFS<sup>c</sup> |  |
| Forest type (dense, open) | Vector25 |  |
| Forest edge (inner and outer) | Calculated | Calculated from the vector25 forest layer |
| Distance to rivers/creeks and lakes | Vector25 |  |
| Proportion of rivers/creeks | Vector25 |  |
| Distance to skilifts and cableways | Vector25 |  |
| Number of sheep & goats | Agricultural statistics | Calculated per pasture-area and community |
| Proportion of federal game reserves | FOEN<sup>d</sup> | Federal hunting reserves with integral or partial protection |
| **Climate** |  |  |
| Mean precipitation in winter | Worldclim | December to February |
| Mean temperature in winter | Worldclim | December to February |

<sup>a</sup> DEM: Digital Elevation Model *swissALTI<sup>3D</sup>*, produced by Federal Office for Topography *Swisstopo*
<sup>b</sup> Vector25: Digital vector map produced by Federal Office for Topography *Swisstopo*
<sup>c</sup> BFS: Swiss Federal Office of Statistics
<sup>d</sup> FOEN: Swiss Federal Office for the Environment

#### Snow conditions for modelling ungulate detection probability

We modelled the detection probability of ungulates as a function of the survey conditions during the transect walks and used several covariates of snow status (Table 3). Snow depth, snow condition and the percentage of snow cover in open and forested areas were recorded directly during snow tracking. In addition, the daily amount of fresh snow at all weather stations in the study regions (56 and 30 in Valais and in the Pre-Alps, respectively) was obtained from the Swiss Institute for Snow and Avalanche Research (WSL/SLF). From these data, we calculated the number of days elapsed between the last snowfall and the date of the transect survey (days since last snowfall, Table 3), assigning each transect to the nearest weather station. We also transformed this continuous variable into a categorical variable describing the snow age. Finally, we recorded the amount of fresh snow that had fallen on the last day with snowfall.

**Table 3.**
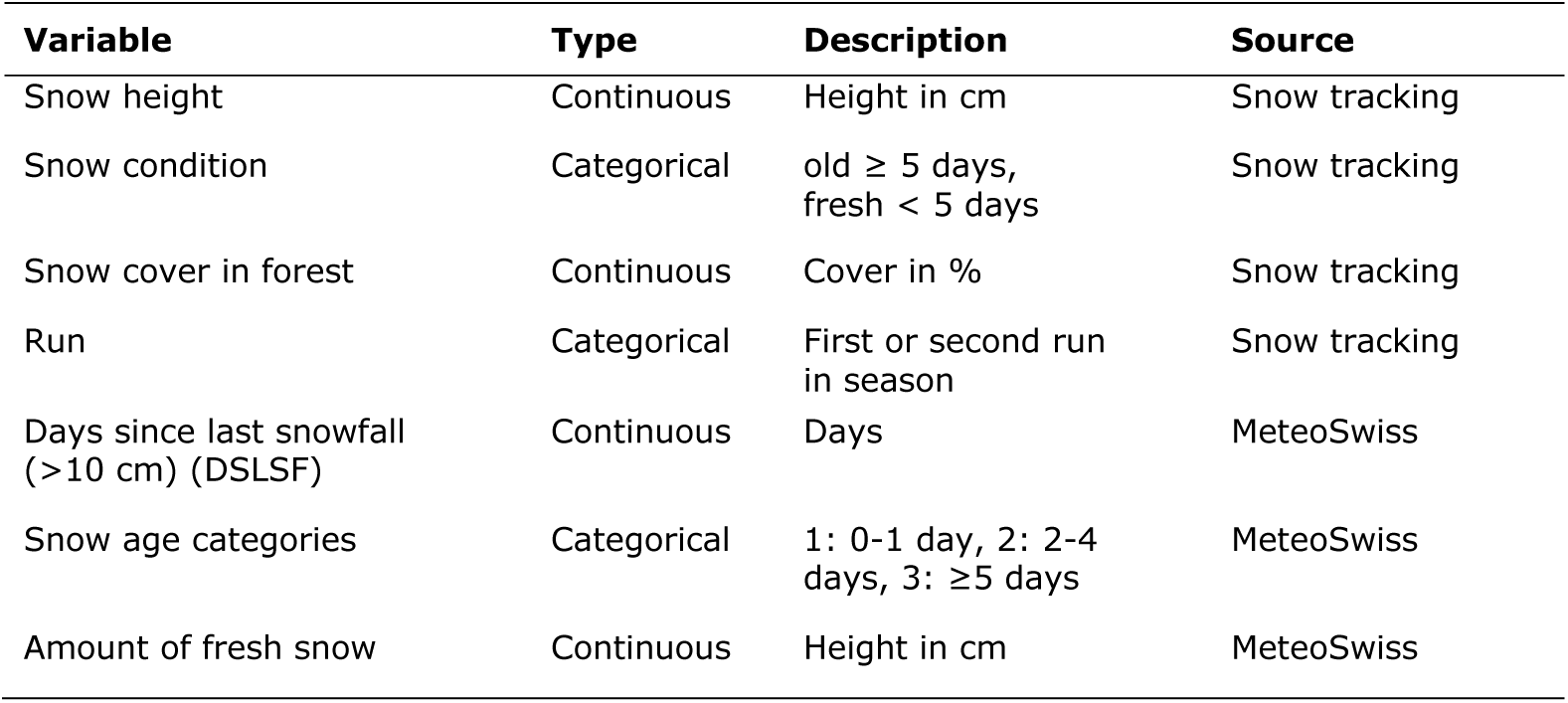
Snow variables used for predicting the detection probability of ungulates based on snowtracking survey data. Variables were recorded directly in the field or derived from the Swiss weather station network MeteoSwiss.

### Surveys for illegal lynx traps

The serendipitous discovery of active lynx traps during the winter 2011/2012 triggered an intensive search of the whole Valais study area in the subsequent years to evaluate the magnitude of this poaching activity. Several traps were operational until 2015. Such contemporary traps had never ever been observed outside Valais, neither in the Pre-Alps nor elsewhere in Switzerland, despite decades of lynx research throughout Switzerland, notably by KORA. Our search consisted of systematically inspecting trees along the 313 1-km long transects (218 in Valais and 95 in the Pre-Alps) for evidence of current and recent historical lynx trap installation (Fig. 2). In effect, lynx trapping systems deployed in Valais are mostly snares typically placed in an obligate passage, i.e. in a bottleneck of microtopographic configurations, such as between big boulders or tall trees, or next to a rock cliff. There, a unique possible path is created by accumulating tree branches in the surroundings so as to form a sort of funnel that channels lynx movement (Fig. 3). A quasi-circular snare is placed vertically at lynx head height, constituting the only possible way through the vegetation, forcing the lynx to pass its head through the snare. When the trap is armed, the snare is connected via a long wire to a large stone that is positioned 2 m above the ground along a tree trunk. Between the snare and the stone, on the trunk surface, there is a hook (often a simple, but thick, folded nail) through which the wire passes. This piece is positioned much higher than the stone itself, which therefore puts the whole system under tension when armed. When a lynx enters its head into the snare and then pulls at it, it triggers the system, which ejects the individual into the air and strangles it (see the contorted lynx heads in Fig. S1). Our quest focused principally on the presence of folded nails, hooks, pulleys and wires on tree trunks, which are often the only parts of a trap system that occur at inactive trapping sites.

**Fig. 3.**
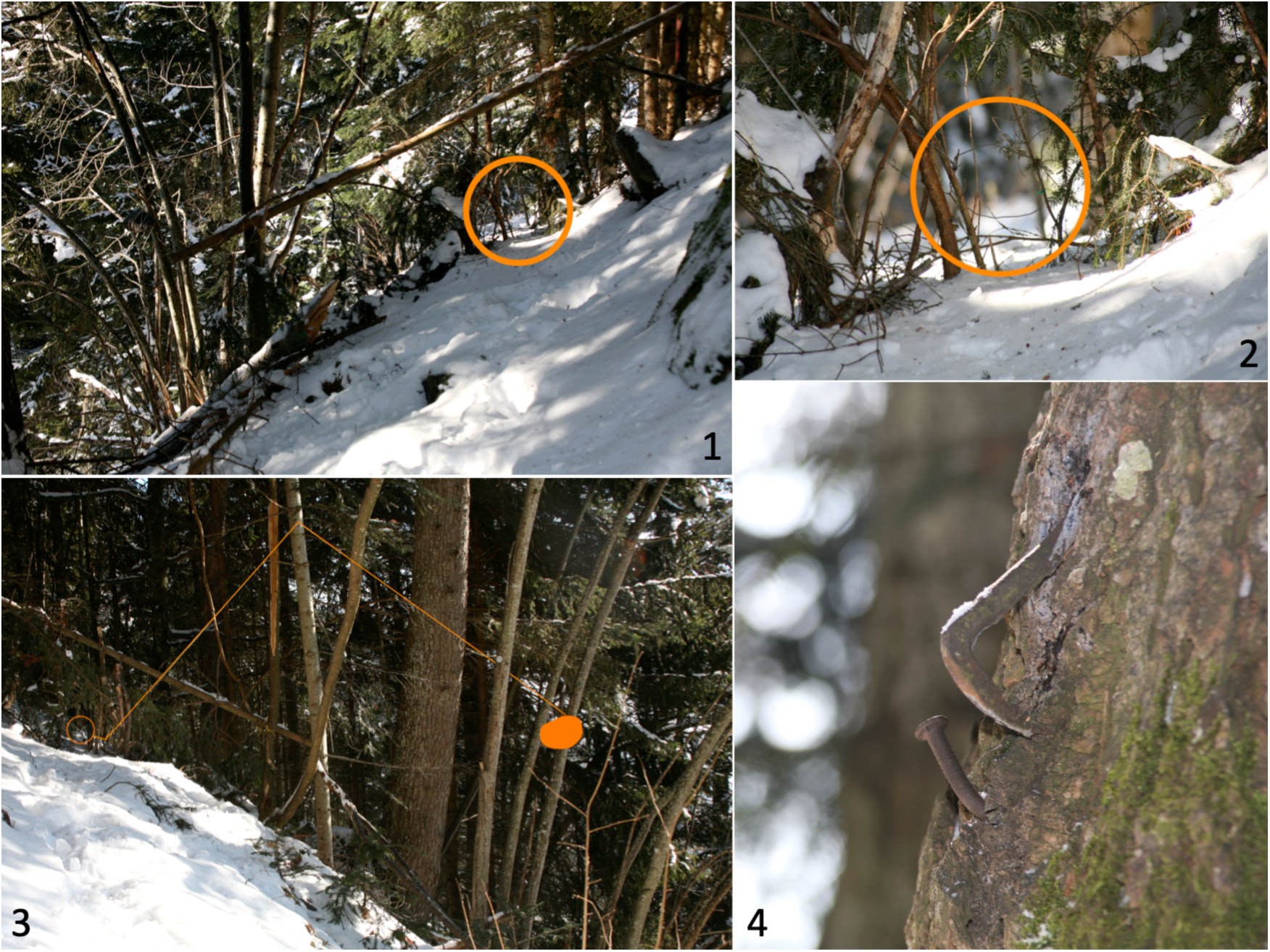
Active snare-ejecting-strangling trap discovered in 2015 in the lynx entrance corridor into Valais from the NW Swiss Alps. 1) The illegal capture system has been installed in an obligate passage along a wildlife trail; branches have been added in the surroundings to augment the channeling effect, i.e. to steer the lynx towards the snare (a bike brake wire) located within the orange circle; 2) a close-up view of the snare itself (cable visible within the orange circle); 3) the trapping system viewed from the other side: the position of the snare is again indicated by the orange circle; the cable bending the system goes through a hook positioned high on the thin tree trunk and goes down towards the right; the system is put under tension with a big stone (highlighted in orange on the right); 4) at an inactive trapping site, sometimes only the middle hook positioned high above the ground is visible. A video showing another active system can be obtained from the authors upon request.

### Testing our hypotheses about the causes of extremely low lynx density in the Valais

#### Hypothesis 1: effect of camera density on lynx densities

To compare trapping efficiency with respect to trail camera density, we compared the number of individual lynx identified by the two survey operations of KORA and Bern University, both conducted in the northwestern part of the Valais study region in the same winter 2013/2014 ^20^. To identify individuals and avoid double-counting, all recorded lynx pictures were compared with each other, as well as with the photographs of all identified lynx stored in the KORA database. Individuals could be identified in all cases through individual fur patterns, even when only one flank of a lynx was pictured. The number of identified lynx was then related to the total surface of suitable lynx habitat in the area, as modelled by Zimmermann ^34^, to estimate lynx density per 100 km^2^. Due to the high density of camera traps and the long duration of surveys we assumed that detection probability was perfect and therefore did not employ (spatial) capture-recapture methods for population assessments.

#### Hypothesis 2: differences in environmental conditions at camera locations

In our comparison of the environmental conditions prevailing at camera locations in normal (Pre-Alps) vs extremely low (Valais) lynx density areas, we first tested for differences in topographic and vegetation conditions (Table 1) using a Wilcoxon rank sum test. Second, we compared topographic and vegetation conditions between camera-traps with and without lynx events to identify the conditions that affected lynx detection in the two study regions. For that analysis, we used the data from two winters (2013/2014 and 2015/2016) for the Pre-Alps whereas for Valais we included the data from three consecutive winters (2013/2014, 2014/2015 and 2015/2016). This choice was based on the fact that camera-traps with lynx captures in Valais were much scarcer than in the Pre-Alps. Each camera with at least one lynx detection within this period was classified as “detection” (“non-detection” otherwise), resulting in 26 out of 102 camera sites with lynx detections (25%) in Valais, compared with 56 cameras out of 79 (71%) in the Pre-Alps.

We then modelled the probability of lynx detection as a function of environmental conditions at the camera sites using generalized linear mixed models (GLMM, package *lme4*” ^35^) with the study region as random factor.

In a first step, we tested for correlations between explanatory variables using the Spearman’s rank correlation coefficient. Of pairs or groups of correlated variables (|r_s_|>0.5) only the one with the better fit in univariate models (according to Akaike’s Information Criterion AIC ^36^) was retained for subsequent analyses. Variables that were significant in a univariate analysis were then included into a multivariate analysis. To identify the most parsimonious model, we tested all possible variable combinations using the dredge function in the R ^37^ package *MuMIn* ^38^and selected the AIC-best model.

#### Hypothesis 3: differences in major ungulate prey densities

To obtain detection-corrected abundance estimates of roe deer and chamois along the transects, N-mixture models were fit with R package *unmarked* ^39^, taking into account the sampling occasion (i.e. first or second survey as described in ^40^) and the snow condition variables as detection variables (Table 3) and the environmental predictors (Table 2) as habitat variables ^40^. For roe deer abundance, we used Poisson models, while for chamois we selected a zero-inflated Poisson distribution. We started calibrating the models using only the detection variables. First, among pairs or groups of correlated variables (Spearman’s |r_s_|>0.5) significantly influencing detection probability, we chose the one that performed best in univariate models, according to AIC corrected for small sample sizes ^41^. Then, models including all possible combinations of variables significantly affecting detection probability were tested and ranked according to their AICc using the *dredge* function of the R-package *MuMIn* ^38^.

Once the best variable combination for detection probability was identified, we additionally included habitat variables into subsequent modelling, adopting the same procedure as above. The best model for each species was then used to predict relative densities (minimum number of individuals per 50 ha) across the whole study area. For this purpose, the proportion (boolean variables) or average (continuous variables) of the relevant environmental predictors (Table 2) were calculated within circular moving windows of 50 ha (radius of 399 m) around each pixel of the study area. We restricted the interpolations to the areas within the same altitudinal range (500–2000 m a.s.l.) and with a similar amount of forest cover (≥14.1%) as found in the 200 m buffer zones around the transects. Additionally, we predicted overall ungulate abundance (N/50 ha) and overall prey biomass (kg/50 ha), assuming average species body masses as reported by the Valais hunting statistics (https://www.vs.ch): 17.5 kg for roe deer (years 2012–2015) and 21.5 kg for chamois (year 2005; both after evisceration).

To compare prey densities between the two study areas, a regular grid of 1-km side length, based on the Swiss grid coordinate system, was applied to the whole study area. For each grid cell, the average density of roe deer, chamois, both prey species pooled, as well as the average prey biomass, were calculated. Then, the densities per cell in both regions were compared using a Wilcoxon rank sum test ^42^. All statistical analyses were performed in R version 3.3.4 ^37^.

#### Hypothesis 4: snare-trap poaching

All signs of recent historical and contemporary presence of lynx snare-ejecting-strangling traps encountered along our 313 1-km survey transects were documented with precise GPS coordinates and photographs. In the area where evidence of illegal trapping systems was found, we intensified the search for other such device locations by walking the majority of trails around the spot, progressing away from the trapping site until no hints for further trapping sites where found over a maximum of 2 km along any such trail.

## Results

### Hypothesis 1: effect of camera density on estimated lynx densities

The two independent surveys carried out during the winter 2013/2014 by KORA and Bern University identified exactly the same seven lynx individuals. The estimated lynx densities obtained were very similar at 0.92 and 0.97 individuals per 100 km^2^ of suitable habitat by the KORA and Bern University, respectively. There was thus no noticeable effect of camera trapping sites density on the estimates of lynx density: this rejects hypothesis 1. The large differences in lynx density between the Pre-Alps and Valais can therefore not be explained by methodological differences in the survey schemes.

### Hypothesis 2: differences in environmental conditions at camera locations

The camera locations at the Valais study sites were characterized by steeper slopes, both near the camera-trap (site scale: Valais: 42.6°; Pre-Alps: 37.2°; W = 3073, p = 0.006) and in the wider surroundings (landscape scale: Valais: 30.0°; Pre-Alps: 26.9°; W = 2656, p < 0.001), by higher proportions of rocks (Valais: 9.8%; Pre-Alps: 3.0%; W = 3104, p = 0.003) and overall obstacles and barriers (Valais: 42.3%; Pre-Alps: 31.6%; W = 2812, p < 0.001) at the site scale, as well as by a higher forest cover (Valais: 62.1%; Pre-Alps: 55.7%; W = 3129, p = 0.01) and density of forest roads (Valais: 6.9 m/km^2^; Pre-Alps: 5.7 m/km^2^; W = 2826, p < 0.001) at the landscape scale. Lynx detection probability at a camera site was only correlated with slope at the landscape scale, with steeper terrain associated with a higher detection probability (GLMM, intercept – estimate = −3.536; SE = 1.683, p = 0.0357; slope = 0.136, SE = 0.041, p < 0.001). These results reveal that the expected detection probability should actually be higher in Valais than in the Pre-Alps. This refutes the hypothesis that camera trapping settings could be the reason for a lower lynx density observed in Valais compared with the Pre-Alps.

### Hypothesis 3: differences in ungulate prey densities

The average per-individual detection probability for roe deer during snow tracking was 49%. It peaked when the snow was less than 5 days-old and fresh snow was not too deep. Roe deer abundance was best explained by an intermediate forest cover, a low winter precipitation and the vicinity of streams and creeks (Table 4a). Detection-corrected prediction of relative (minimum) roe deer abundance at the transect sites ranged from 0.23 to 11.11 (mean: 3.81) individuals. Predicted relative roe deer densities across both study regions ranged between 0.44 to 6.61 individuals per 50 ha (Fig. 4a). For chamois, mean per-individual detection probability was 30%. The depth of fresh snow had a negative, but non-significant effect on detection probability. Chamois abundance was positively influenced by slope, intermediate grassland cover and by distance to lakes (Table 4b). Detection-corrected predictions of abundance along the transects ranged from 0 to 10.03 (mean: 1.41) individuals, while predicted relative chamois densities in the two study regions ranged from 0 to 37.8 individuals per 50 ha (Fig. 4b). Overall prey densities for both species combined ranged from 0.5 to 42.6 individuals per 50 ha, total biomass per 50 ha from 8.4 to 896.4 kg (Fig. 4c, d). Overall relative prey densities differed between the two study regions. With an average of 2.41 individuals per 50 ha in the Pre-Alps, compared to 3.29 per 50 ha in Valais (Wilcoxon-test: W = 679280, p < 0.001), roe deer densities were significantly higher in Valais (Fig. 5a). A similar pattern existed for relative chamois densities (Wilcoxon-test: W = 888420, p < 0.001), with an average of 3.44 individuals per 50 ha in the Pre-Alps and 4.75 individuals per 50 ha in Valais (Fig. 5b). Total prey density (Pre-Alps: 5.85; Valais: 8.04) and biomass per 50 ha (Pre-Alps: 116.06 kg; Valais: 159.61 kg) were also substantially higher in Valais (Wilcoxon-test: W = 764310, p < 0.001; W = 779900, p < 0.001, respectively) (Fig. 5c, d). Thus, the third hypothesis, that prey supply for lynx may be lower in Valais than in the Pre-Alps, which could explain a lower lynx density in Valais, was convincingly rejected.

**Table 4.** Best N-mixture models for predicting relative densities of roe deer (a) and chamois (b). Estimates, standard error (SE) and p-values are given for both the abundance and detection analysis.

| Variable | Estimate | SE | p-value |
| --- | --- | --- | --- |
| **(a) Roe deer** |  |  |  |
| Abundance (log-scale) |  |  |  |
| Intercept | 0.625 | 0.334 | 0.062 |
| Forest | 4.644 | 0.976 | <0.001 |
| Forest squared | -3.583 | 0.862 | <0.001 |
| Precipitation winter | -0.006 | 0.002 | <0.001 |
| Rivers and Creeks | -2.166 | 0.777 | 0.005 |
| Detection (logit-scale) |  |  |  |
| Intercept | 0.260 | 0.330 | 0.431 |
| Snow Condition | -0.337 | 0.130 | 0.010 |
| Run | -0.142 | 0.089 | 0.111 |
| Height of fresh snow | -0.012 | 0.005 | 0.016 |
| **(b) Chamois** |  |  |  |
| Abundance (log-scale) |  |  |  |
| Intercept | -1.402 | 0.769 | 0.068 |
| Slope | 0.068 | 0.018 | <0.001 |
| Grasslands | 2.505 | 1.230 | 0.042 |
| Grasslands squared | -3.280 | 1.628 | 0.044 |
| Forest edges | 0.024 | 0.014 | 0.075 |
| Distance to lakes | 0.175 | 0.066 | 0.008 |
| Sheep and goats | -3.428 | 2.188 | 0.117 |
| Detection (logit-scale) |  |  |  |
| Intercept | -0.482 | 0.464 | 0.300 |
| Snow height | -8.86e-4 | 0.004 | 0.843 |
| Height of fresh snow | -0.016 | 0.009 | 0.077 |

**Table 5.** History of lynx poaching with snare-ejecting-strangling traps in Valais, southwestern Switzerland, based on media broadcast and justice decisions. X is the hunter eventually convicted in 2015 for the use of illegal trapping devices.

| Year | Event and source | Procedure | Administration | Outcome |
| --- | --- | --- | --- | --- |
| 1995 | X portrayed with a rifle and two lynx (Fig. S1) <sup>14, 19</sup> | Anonymous sending of the picture to the Swiss Confederation and broadcast of the picture in mass media. The head of the federal wildlife service said it had sent the picture for denunciation to the local court. | Local court (Bas-Valais). | Case dismissed by court because it said it never received the picture from the federal agent. |
| 2005 | A hiker is caught in a snare trap by his foot and has difficulties to free himself | Complaint filed by the victim at the local police; visit of the trapping system with a game warden. Complaint after a few months to the same game warden that the trap was again active. The trap remained on site until 2015. | Local police and local court (Bas Valais) | Trap removed during a few months, but then re-installed (active until 2015 when we discovered and dismantled it)<br>Case dismissed by court for unknown reasons. |
| 2013 | X posing in mass media, claiming to have trapped to death 10 lynx with snares | Reports to the cantonal wildlife and game service by WWF and Pro Natura (NGOs) | Local police, game wardens and local court (Bas Valais) | Case dismissed by court after it ruled the inquiry by game wardens and local police could not prove the existence of trapping systems.<br>Case dismissed by court: X claimed the two lynx he was portrayed with in 1995 were found crashed by vehicle (one killed by himself while driving). His statement was backed by a local game warden. |
| 2015 | Discovery of 17 snare-ejecting-strangling traps by paper authors; 3 were active | Dismantling of traps and delivery of materials to cantonal criminal police after consulting Attorney General; DNA investigations | Cantonal court (Valais), i.e. higher jurisdiction than previously. | Court convicts X for use of illegal hunting methods. |

**Figure 4.**
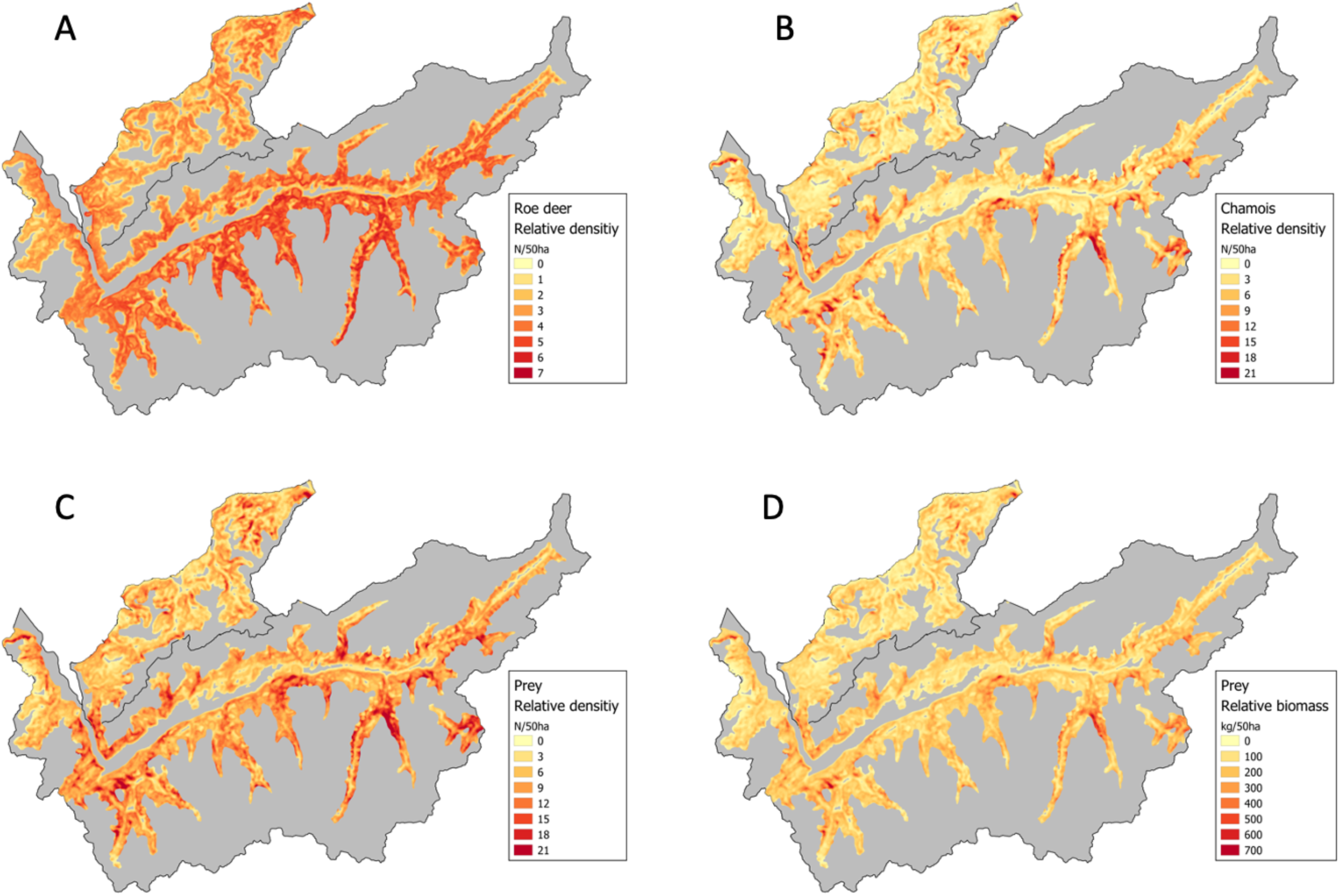
Predicted relative densities of A) roe deer (ranging from 0.44 to 6.61 individuals per 50ha); B) chamois (0 to 37.76 per 50 ha); C) all ungulate prey together (0.47 to 42.59 per 50 ha) and D) predicted relative ungulate biomass (8.38 to 896.39 kg per 50 ha). Predictions were only made for the yellow to red coloured area.

**Figure 5.**
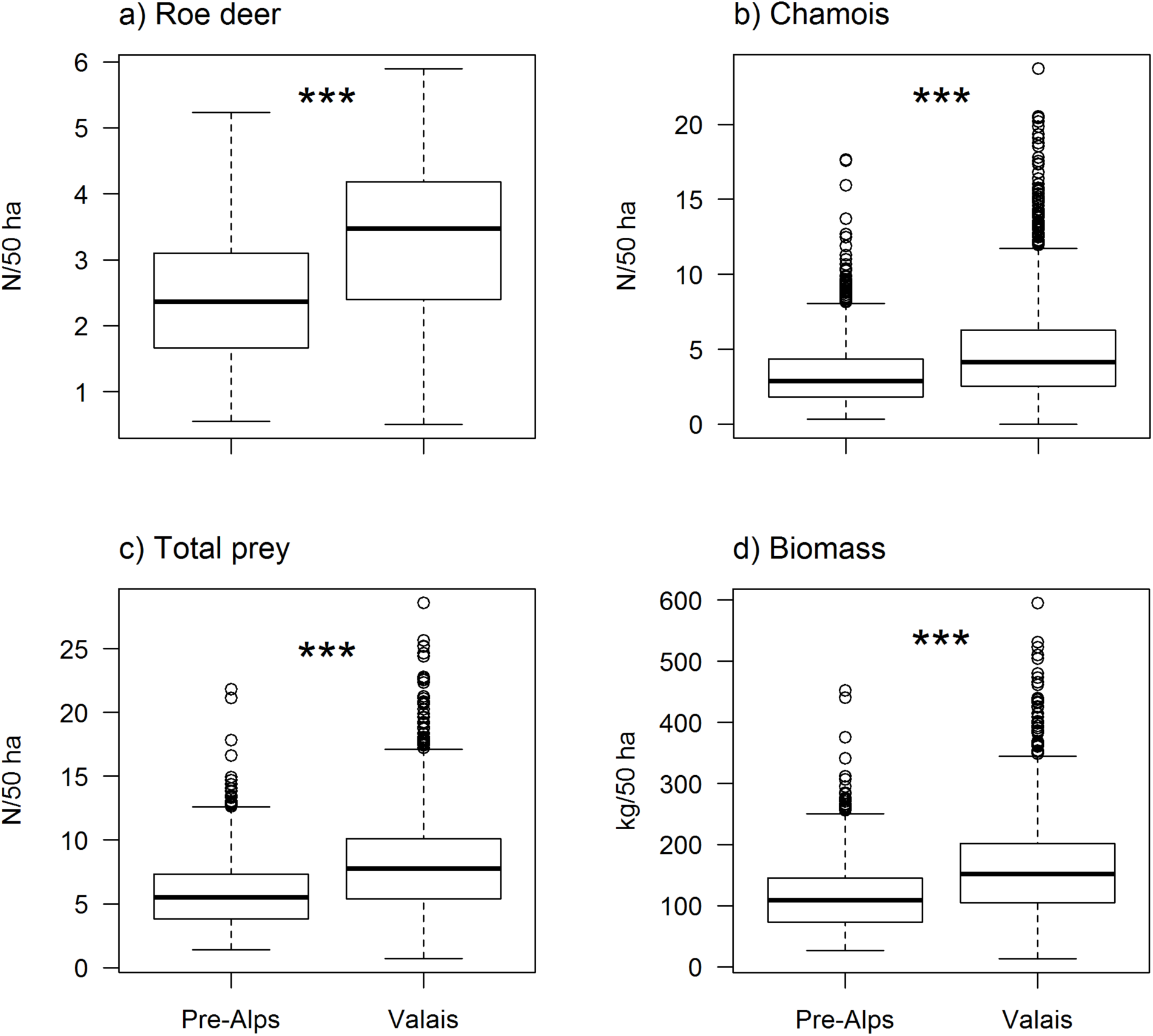
Comparison of ungulate densities and biomass between the NW Pre-Alps and Valais: A) roe deer; B) chamois; C) total prey density and D) biomass, in number of individuals and kg per 50 ha, respectively.

**Figure 6.**
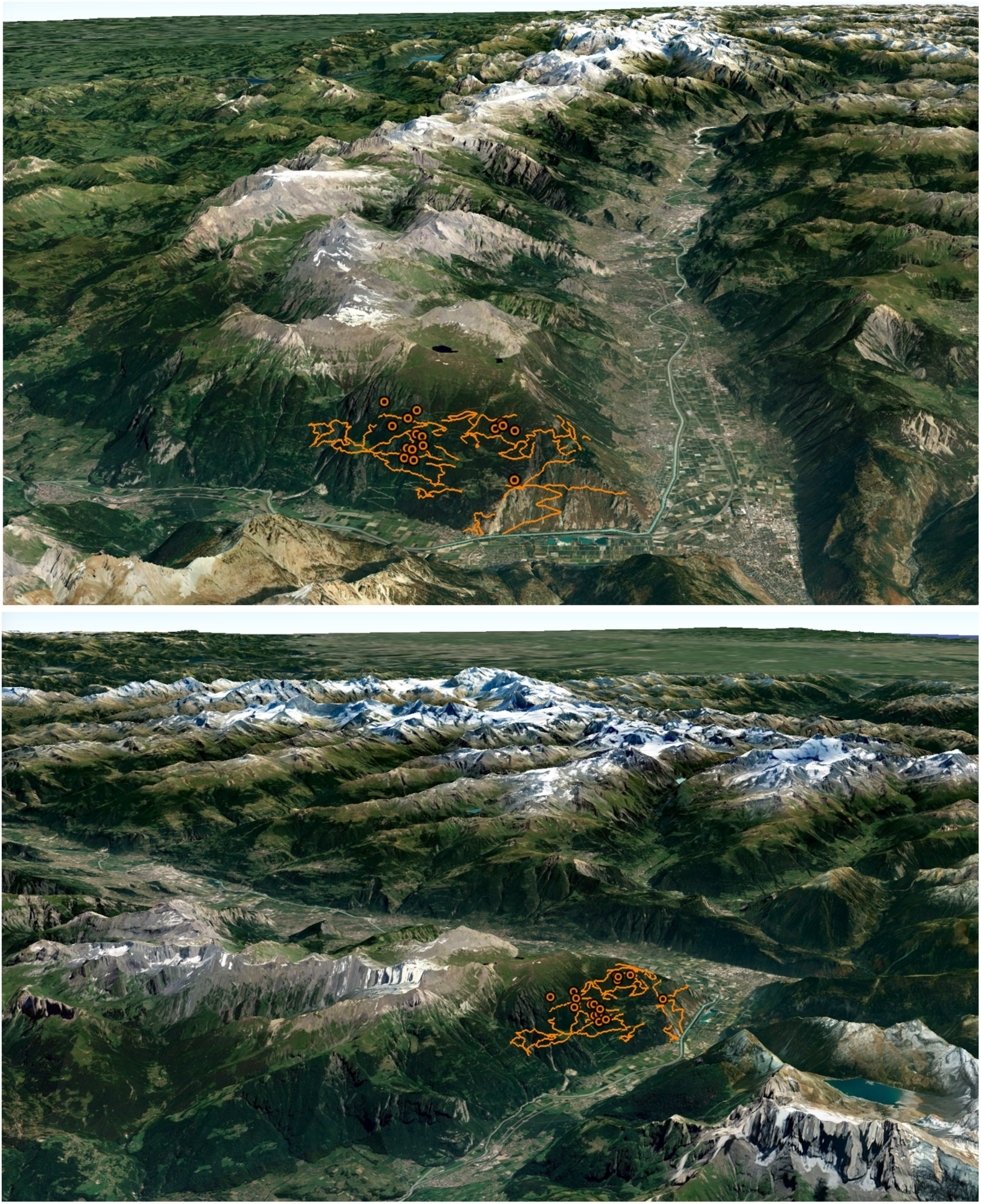
Google Earth ™ views (from the SW, upper projection; from the NW, lower projection; compare to Fig. 1) of the entrance corridor into Valais, from the lynx-populated NW Alps, showing the 39 km of trails (orange lines) explored for traps as well as the 17 snare-strangling-ejecting trap sites (orange dots) discovered (two traps are so close that they appear as a simple dot). The forest belt as well as the barriers to dispersal constituted by the high mountain chains are obvious.

### Hypothesis 4: trap poaching

We found evidence for current and recent past use of snare-ejecting-strangling lynx traps only in a key area surrounding the village of Champex d’Alesses sur Dorénaz (postcode 1905). A more in-depth prospection along 39 km of trails over a total area of 15 km^2^ around the village enabled us to discover the presence of 17 different poaching trap sites in 2015, with evidence of past or present installation of snare-ejection-strangling lynx traps (Fig. 3 & 6). The 17 trapping systems were concentrated in an area of ca 4 km^2^, thus at a density of ca 4 traps per km^2^. Three traps were armed the very day when we discovered them, while all the other 14 were currently inactive, with their most-recent years of deployment unknown. We dismantled all active traps and delivered them to the cantonal police. In contrast, no signs of similar trap systems were discovered along any of the other 1-km transects in the two study areas, suggesting that snare-trapping was a fairly local activity around Champex d’Alesses. However, the network of traps covered exactly the single main immigration corridor for lynx into Valais from the North, i.e. from the only healthy lynx population adjacent to the Valais from where immigration might reasonably take place: the high-density, thriving lynx populations in the Pre-Alps (Fig. 1). It appears likely that the strategic location of this “defence of Valais” against immigrating lynx may have leveraged its substantial impact by greatly reducing the immigration of lynx into Valais. We therefore failed to reject the hypothesis that poaching reduced lynx densities in Valais.

## Discussion

Our study aimed at elucidating the reason for why currently the lynx density in Valais is only about 20% of the expected density, with an almost complete absence of reproduction for many years ^20, 27^. Our analysis rejected all methodological and biological hypotheses but provided evidence in support of an intense local and strategically placed trapping pressure, offering a plausible explanation for the current restricted distribution range and extremely low density of lynx in Valais ^20, 27^. Note that if our study did not find any evidence for other snare traps elsewhere in Valais despite intensive search, it did not investigate the role of further possible poaching using other methods throughout the study area.

First, we found that a density of camera-traps 36% lower in Valais compared to the Pre-Alps did not materially affect the probability of detection of individual lynx. This means that the two monitoring schemes operated in the Pre-Alps and Valais, by KORA and Bern University, respectively, converged in their estimates of lynx population size ^20^. Second, we found that the environmental conditions surrounding the camera-trap sites (which were preferentially placed at “obligate passages”) should have made it even more likely for lynx to be photo-captured in Valais than in the Pre-Alps. This suggests that the probability of detecting the presence of a lynx in Valais should be higher in Valais than in the Pre-Alps. Third, the density and total biomass of the two key prey species, roe deer and chamois, were substantially higher in Valais than in the Pre-Alps, indicating that a reduced prey base cannot explain the much lower lynx population density in Valais ^43^. In contrast, the fourth hypothesis, poaching, was supported by the discovery of a dense network of snare-ejecting-strangling lynx traps in the principal immigration corridor into the Upper Rhône valley (Valais) from the dense populations occurring in the Pre-Alps to the North. Poaching of lynx thus remains as the only plausible explanation for the drastic population limitation of lynx in Valais.

Forest corridors are essential for lynx dispersal. Contrary to other top carnivores such as wolf and bear (*Ursus arctos*), which are regularly observed in open alpine habitat well above the timberline, lynx are typical forest-dwellers, even during long-distance dispersal. There is only one radio-documented case of a lynx dispersing into Valais directly from the North over a high Alpine pass, out of several dozen of radio-tracked dispersal routes ^20^. The almost exclusive possibility for lynx to enter Valais is therefore through the aforementioned forested corridor (Fig. 1) through which most individuals emigrate from the high-density Pre-Alpine populations into Valais^20^. It is precisely in this unique narrow and steep wooded corridor where the dense network of illegal traps was located. With a very high density of trap systems of 4 per km^2^, it appears highly likely that this illegal trapping effort could have a major effect on lynx population dynamics in the entire Valais ^44, 45^, although the precise magnitude of its impact remains of course unknown.

Biollaz et al. ^20^ established the existence of a gradual spatial decrease in lynx density from that entry corridor in the West towards the East and South of Valais (Fig. 2 in ^20^), with some of the main tributaries of the Pennine Alps (South of the Rhône), such as the valleys of Hérens and Anniviers, being almost devoid of any lynx today ^20^. This has recently been corroborated by further camera-trap surveys carried out by KORA ^27^ and contrasts sharply with the situation in the 1980s, when the species occurred over a much wider area of Valais ^13^, notably South of the river Rhône and within the aforementioned valleys. The question that arises then is whether the mortality generated by the unveiled local trap network could have contributed to reduce the overall Valais lynx population to the extremely low density observed at present, which contrasts so strongly with the projections of the habitat suitability model by Zimmermann ^34^ and with expectations based on the highly abundant prey supply, as evidenced in the present study. It is fairly likely that some level of poaching also occurs in other parts of Valais as well, but relying on methods other than snare traps as we found snares only in the main immigration corridor into Valais, this despite intensive search all over. Some Swiss hunters publicly claim to regulate lynx because they believe hunting is threatened by the return of top predators ^46-48^ and there is evidence from mounted taxidermized lynx that illegal hunting actually did occur in Valais ^14^. Our interpretation of the above-mentioned density gradient is that poaching, possibly using other methods that snare traps (i.e. guns), is taking place throughout Valais at some unknown level, with lynx steadily trying to colonize the Rhône valley from the thriving, better protected populations in the Pre-Alps. A fraction of these potential immigrants was continuously removed by the trap network situated in the main entry corridor ^44^, impeding immigration to compensate for the probable poaching-related losses within Valais. Hence, if lynx trapping may possibly not be the only cause of the extremely low lynx population density at present, it certainly did contribute to it.

A remarkable aspect of the illegal poaching activity using snares is that it had been officially known since at least 1995 by the cantonal game wardens and administrative authorities, as assessed by the NGOs Pro Natura ^48^ and WWF (WWF Valais, *in litt*.) and even by Swiss national television ^49^. In that year, a licensed Valais hunter (hereafter X) had been portrayed, not far from his place of residence and the trap network we discovered, proudly staging with a gun and two dead lynx, which had obviously been killed by strangulation, based on the contorted posture of their heads (Fig. S1; see also ^14^). The picture was sent anonymously to the federal wildlife service who reported to have forwarded it to the Valais judiciary. In 2013, X claimed in an interview to a Swiss magazine that he had “*trapped to death 10 lynx with snares*” ^44^, thus corroborating our hypothesis of a possible population-level impact of the unveiled trap network. Following this broadcast, the Game and fishery service of Valais, together with WWF and Pro Natura, filed a charge against X, but the court dismissed the case in 2014, arguing that the investigations by the local police and game wardens could not provide evidence of the existence of any operational traps while the hunter pretended that the picture of 1995 was staged, with the two lynx being roadkills found dead (WWF Valais, *in litt*.). One may question the quality of the inquiry in 2014 given that in 2015 we discovered the network of 17 snare-ejecting-strangling systems, none being of recent installation. Among them, three were armed, i.e. active in operation. We dismantled the traps and delivered them to the cantonal criminal police, i.e. to the highest regional judicial authority, while filing a formal complaint against X who was eventually convicted for use of illegal hunting devices after his DNA was found on the traps (Fig. 3) ^45^.

To understand the broader context, it is worth mentioning that in 2005 a hiker was caught by the foot in one of these snare traps ^49^. He filed a lawsuit at a local police station and was accompanied by a cantonal game warden who took pictures of the trapping system at the site. Although this resulted in the temporary removal of the trap, the victim found the same trap operational again only a few months later, where it remained *in situ* until we dismantled it in 2015. This suggests that the same trap network may have been operating for years, if not decades. It was clear that the intervention of local game wardens was not sufficient to stop these illegal activities during at least a full decade.

This study, as well as the above and other similar facts reported by both the media and the judiciary since 1995 supports the hypothesis that lynx poaching had long taken place in Valais, and that – if strategically placed – the illegal activities of perhaps a single person may represent a serious threat to a small and highly isolated population ^20^. Obviously, lynx trapping continued unabated until we approached the cantonal Attorney General in 2015. From that very moment, it was the cantonal criminal police, instead of local law-enforcement agents, that was mandated for conducting the inquiries about carnivore trap-poaching ^45^. This seems to have triggered a series of serious investigations (including the decisive use of DNA analysis) for the first time, which fairly rapidly led to a conviction ^45^.

Poaching in populations of large carnivores may be a source of additive mortality that can have severe consequences, especially for small populations and even more so when they are highly isolated. In Valais, poaching is likely to represent a threat to lynx: vast zones that had been recolonised as early as the 1980s are now devoid of this felid ^13, 20^. Other causes (e.g. diseases or inbreeding depression) can be ruled out in our opinion because other Swiss populations would then have showed signs of such impacts on demography as well. In light of all the evidence presented in our study, long-lasting poaching, using traps and possibly other methods as well, appears to be the most plausible cause for the extremely low lynx density found in Valais today. In Valais, poaching of legally protected species has also been confirmed for bearded vulture (*Gypaetus barbatus*) and wolf, with recurrent events for the latter species in the recent times, indicating a certain “frontier mentality” and a general disrespect for biodiversity conservation law at least among some people in this geographically isolated mountain region in the Swiss Alps. The history of confirmed, but unaddressed, poaching events for years renders legitimate raising the question of whether some level of proximity between poachers and local law enforcement agents may have allowed poaching to keep continuing unabated. By not taking sufficient measures against the snare-traps deployment, even after a person was injured, local law enforcement authorities appear to have sent a signal of tacit approval to lynx poaching activity. This alone may explain why the lynx trapper even felt emboldened to appear in the mass media with a belief of total impunity ^44^.

Cases of collusion between game wardens, police forces and poachers are known from other European countries. In Norway in 2016, a police chief inspector conducting investigations on wolf poaching sent text messages to an accused hunter, explaining him how to answer questions from the criminal police so as to have the case dismissed ^50^. This case reached the Supreme Court of Norway, which ruled in 2018 that the breach of his service obligations was serious because his advice to the poacher could reduce the trust among the public regarding the ability and willingness of the police to investigate criminal matters ^50^. Also in Norway, in 2015, a retired policeman was found guilty of having helped concealing evidence of illegal wolf hunting ^51^. It is worth noting that this occurred in a country that is comparable to Switzerland ^52^ in consistently scoring among the top nations in the world for following the rule of law, with a quasi-absence of corruption. Informal discussions we have had with other large carnivore researchers in northern Europe indicate that informing local police about large carnivore poaching may indeed merely amount to informing poachers about the visibility of their illegal activity. In effect, in many instances, local police forces seem to treat poaching, especially of large carnivorous predators, as being a socially acceptable crime. It is clear that such a mentality will be a serious impediment to proper investigations.

In the Swiss context, poachers and official cantonal game wardens often belong to the same local social network. In effect, in the Swiss federal political system, police forces are mostly appointed locally, at the cantonal level. This is in contrast to France and Italy, for instance, where law enforcement agents in the countryside are usually not appointed in the area they originate from ^53^, but are re-allocated across the country to prevent corruption, which has led to strong charges against lynx poachers ^48^. The belief among poachers that they are immune from prosecution likely emboldens them to continue their activity. If a coherent strategy aimed at reducing poaching of large carnivores needs to be integrative in all respects, from prevention to legal prosecution^54^, the risk of a possible tolerance towards poaching that local game police forces may display must be addressed head-on. In other terms, efforts to control poaching may therefore require to police the police as 2^nd^ century Roman poet Juvenal wrote in his Satires “Quis custodiet ipsos custodes”. In this respect, we would recommend Switzerland and other European countries where poaching is suspected of threatening top predators to investigate it at the highest possible administrative level or establish a centralized police force tasked with investigating major environmental crimes, including the illegal hunting of large carnivores.

## Acknowledgements

We thank all the people who made this study possible, in particular Dr Reini Schnidrig-Petrig, Swiss Federal Office for the Environment, who supported financially our research programme. Particular thanks to the attorney general and the criminal police for their inquiry about the perpetuator of lynx trap installation and damages to our trail camera-trap network.

---

**Figure S1.**
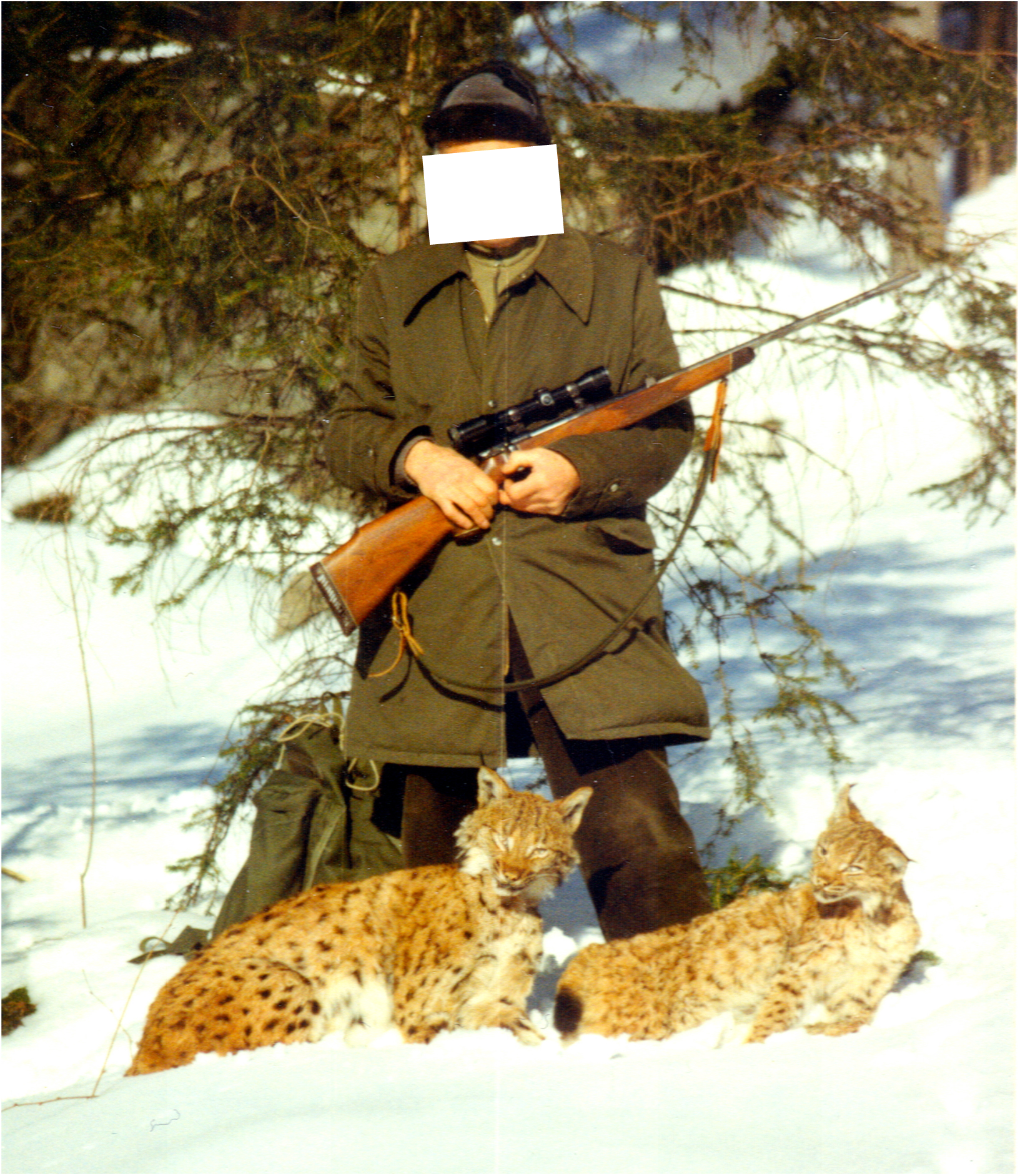
The hunter X, eventually convicted for illegal hunting in 2015, staging with two dead lynx and a gun (by courtesy of ^14^ and the photographer, Jean-Claude Tornay). The contorted position of their heads indicates that the two lynx have been strangled and not shot, which could be assessed by one of the authors while visiting the armory where one of these two lynx, recognizable by its fur pattern, was long exposed stuffed.

**Figure S2.**
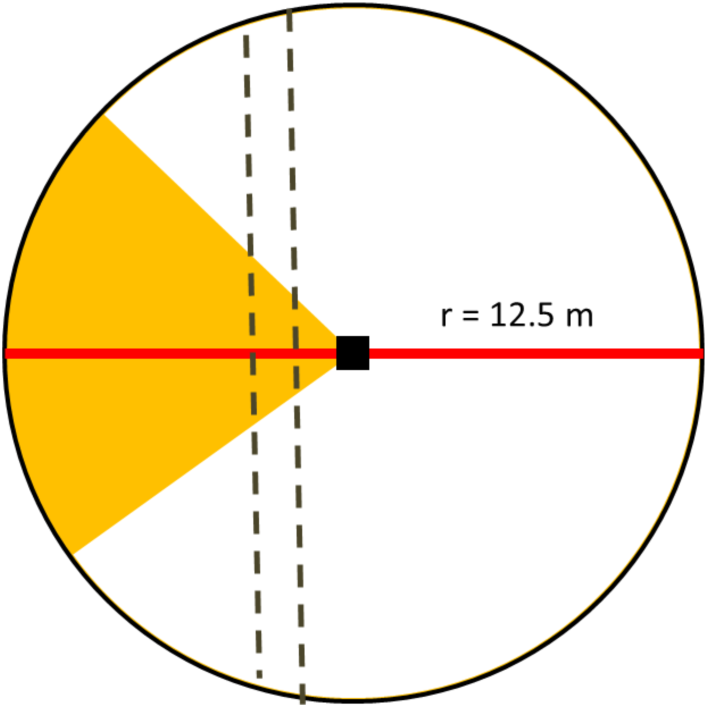
Environmental conditions in the vicinity of a trail camera-trap were assessed either within a 12.5-m radius plot centered on the camera (depicted by a black square) or along a 25-m long transect (red line). The yellow triangle symbolises the approximate camera coverage. The dashed lines depict a dirt road or trail where a lynx is expected to transit.

## Notes

### Competing Interest Statement

The authors have declared no competing interest.

## References

1 Treves, A. & K. U. Karanth. 2003. Human-carnivore conflict and perspectives on carnivore management worldwide. Conservation Biology 17: 1491–1499.

2 Chapron, G. & J. V. López-Bao. 2016. Coexistence with Large Carnivores Informed by Community Ecology. Trends in Ecology & Evolution 31: 578–580.

3 López-Bao, J. V., J. Bruskotter & G. Chapron. 2017. Finding space for large carnivores. Nature Ecology & Evolution 1: 0140.

4 Ripple, W. J., J. A. Estes, R. L. Beschta, C. C. Wilmers, E. G. Ritchie, M. Hebblewhite, J. Berger, B. Elmhagen, M. Letnic, M. P. Nelson, O. J. Schmitz, D. W. Smith, A. D. Wallach & A. J. Wirsing. 2014. Status and Ecological Effects of the World’s Largest Carnivores. Science 343.

5 Chapron, G., P. Kaczensky, J. D. C. Linnell, M. von Arx, D. Huber, H. Andrén, J. V. López-Bao, M. Adamec, F. Álvares, O. Anders, L. Balčiauskas, V. Balys, P. Bedő, F. Bego, J. C. Blanco, U. Breitenmoser, H. Brøseth, L. Bufka, R. Bunikyte, P. Ciucci, A. Dutsov, T. Engleder, C. Fuxjíger, C. Groff, K. Holmala, B. Hoxha, Y. Iliopoulos, O. Ionescu, J. Jeremić, K. Jerina, G. Kluth, F. Knauer, I. Kojola, I. Kos, M. Krofel, J. Kubala, S. Kunovac, J. Kusak, M. Kutal, O. Liberg, A. Majić, P. Mínnil, R. Manz, E. Marboutin, F. Marucco, D. Melovski, K. Mersini, Y. Mertzanis, R. W. Mysłajek, S. Nowak, J. Odden, J. Ozolins, G. Palomero, M. Paunović, J. Persson, H. Potočnik, P.-Y. Quenette, G. Rauer, I. Reinhardt, R. Rigg, A. Ryser, V. Salvatori, T. Skrbinšek, A. Stojanov, J. E. Swenson, L. Szemethy, A. Trajçe, E. Tsingarska-Sedefcheva, M. Váňa, R. Veeroja, P. Wabakken, M. Wölfl, S. Wölfl, F. Zimmermann, D. Zlatanova & L. Boitani. 2014. Recovery of large carnivores in Europe’s modern human-dominated landscapes. Science 346: 1517.

6 Epstein, Y. & G. Chapron. 2018. The Hunting of Strictly Protected Species: The Tapiola Case and the Limits of Derogation under Article 16 of the Habitats Directive. European Energy and Environmental Law Review 27: 78–87.

7 Liberg, O., G. Chapron, P. Wabakken, H. C. Pedersen, N. Thompson Hobbs & H. Sand. 2012. Shoot, shovel and shut up: Cryptic poaching slows restoration of a large carnivore in Europe. Proceedings of the Royal Society B: Biological Sciences 279: 910–915.

8 Andrén, H., J. D. C. Linnell, O. Liberg, R. Andersen, A. Danell, J. Karlsson, J. Odden, P. F. Moa, P. Ahlqvist, T. Kvam, R. Franzén & P. Segerström. 2006. Survival rates and causes of mortality in Eurasian lynx (Lynx lynx) in multi-use landscapes. Biological Conservation 131: 23–32.

9 Sindičić, M., T. Gomerčić, J. Kusak, V. Slijepčević, Đ. Huber & A. Frković. 2016. Mortality in the Eurasian lynx population in Croatia over the course of 40 years. Mammalian Biology 81: 290–294.

10 Heurich, M., J. Schultze-Naumburg, N. Piacenza, N. Magg, J. Červený, T. Engleder, M. Herdtfelder, M. Sladova & S. Kramer-Schadt. 2018. Illegal hunting as a major driver of the source-sink dynamics of a reintroduced lynx population in Central Europe. Biological Conservation 224: 355–365.

11 Červený, J., P. Koubek & L. Bufka. 2002. Eurasian Lynx (Lynx Lynx) and its Chance for Survival in Central Europe: The Case of the Czech Republic. Acta Zoologica Lituanica 12: 428–432.

12 Liberg, O., G. Chapron, P. Wabakken, H. C. Pedersen, N. T. Hobbs & H. Sand. 2012. Shoot, shovel and shut up: cryptic poaching slows restoration of a large carnivore in Europe. Proceedings of the Royal Society. Series B, Biological sciences 279: 910–915 (plus 916 p. supplementary material).

13 Haller, H. 1992. Zur Ökologie des Luchses Lynx lynx im Verlauf seiner Wiederansiedlung in den Walliser Alpen. Verlag Paul Parey, Hamburg. 62 p. pages.

14 Breitenmoser, U. & C. Breitenmoser-Würsten. 2008. Der Luchs: ein Grossraubtier in der Kulturlandschaft. Salm Verlag, Wohlen/Bern. 537 pages.

15 Arlettaz, R., R. Imstepf, A. Jacot, P.-A. Oggier, B. Posse, J.-N. Pradervand, E. Revaz, P. Salzgeber, A. Sierro, B. Wolf, U. Zimmermann & S. Zurbriggen. 2019. Oiseaux et biodiversité du Valais: comment les préserver. Station ornithologique suisse, Sempach. pages.

16 Molinari-Jobin, A., S. Wölfl, E. Marboutin, P. Molinari, M. Wölfl, I. Kos, M. Fasel, I. Koren, C. Fuxjíger, C. Breitenmoser, T. Huber, M. Blažič & U. Breitenmoser. 2012. Monitoring the Lynx in the Alps. Hystrix 23: 49–53.

17 Molinari-Jobin, A., F. Zimmermann, C. Angst, C. Breitenmoser-Würsten, S. Capt & U. Breitenmoser. 2006. Status and distribution of the lynx in the Swiss Alps 2000–2004. Acta Biologica Slovenica 49: 3–11.

18 Molinari-Jobin, A., F. Zimmermann, C. Breitenmoser-Würsten, S. Capt & U. Breitenmoser. 2001. Present status and distribution of the lynx in the Swiss Alps. Hystrix n.s. 12 17–27.

19 Zimmermann, F., A. Molinari-Jobin, A. Ryser, C. Breitenmoser-Würsten, E. Pesenti & U. Breitenmoser. 2011. Status and distribution of the lynx (Lynx lynx) in the Swiss Alps 2005–2009. Acta Biologica Slovenica 54: 74–84.

20 Biollaz, F., S. Mettaz, F. Zimmermann, V. Braunisch & R. Arlettaz. 2016. Statut du Lynx en Valais quatre décennies après son retour: suivi ou moyen de pièges photographiques. Bulletin de la Murithienne 133: 29–44.

21 Molinari, P., L. Rotelli, M. Catello & B. Bassano. 2001. Present status and distribution of the Eurasian lynx (Lynx lynx) in the Italian Alps. Hystrix 12: 3–9.

22 Biollaz, F. & R. Arlettaz. 2017. Le problème du Valais avec les superprédateurs. Le Temps.

23 Fournier, J.-R. 2010. Motion Jean-René Fournier. Révision de l’article 22 de la Convention de Berne. Bulletin officiel du Conseil des Etats, Parlement suisse Motion 10.3264.

24 Gonseth, Y., T. Wohlgemuth, B. Sansonnens & A. Buttler. 2001. Die biogeographischen Regionen der Schweiz. Erlíuterungen und Einteilungsstandard. Page 48. Bundesamt für Umwelt (BAFU), Bern.

25 Zimmermann, F., F. Kunz, K. Rhein, M. Shepherd, P. Tschanz, C. Breitenmoser-Würsten & U. Breitenmoser. 2016. Abondance et densité du lynx dans le Nord-Ouest des Alpes: estimation par capture-recapture photographique dans l’aire de référence étendue au canton de Vaud dans le C-VI durant l’hiver 2015/16. KORA report nb. 74. KORA, Muri.

26 Breitenmoser, U. & H. Haller. 1987. Zur Nahrungsökologie des Luchses (Lynx lynx) in den schweizerischen Nordalpen. Zeitschrift für Síugetierkunde 52: 168–191.

27 Zimmermann, F., J. Küttel, C. Breitenmoser-Würsten, U. Breitenmoser & F. Kunz. 2019. Monitoring déterministe du lynx avec les pièges-photos dans le Sud du Bas-Valais IVd durant l’hiver 2018/19, Bertm. 19 pages.

28 F., Z. & F. D. 2016. Capture-recapture methods for density estimation. Pages 95-141 in R. F. and Z. F., editors. Camera trapping for wildlife research. Pelagic Publishing.,Exeter, UK.

29 Roder, S., F. Biollaz, S. Mettaz, F. Zimmermann, R. Manz, M. Kéry, S. Vignali, L. Fumagalli, R. Arlettaz & V. Braunisch. 2020. Deer density drives habitat use of establishing wolves in the Western European Alps. Journal of Applied Ecology 57: 995–1008.

30 QGIS Development Team. 2017. Geographic Information System. Version 2.18.9 Las Palmas. Open Source Geospatial Foundation Project, http://www.qgis.org.

31 Zimmermann, N. E. & D. W. Roberts. 2001. Final Report of the MLP climate and biophysical mapping project. WSL, Birmensdorf, Switzerland. pages.

32 Hamr, J. 1985. Seasonal home range size and utilisation by female chamois (Rupicapra rupicapra) in Northern Tyrol. Pages 106–116 in S. Lovari, editor. The Biology and Management of Mountain Ungulates. Croom Helm, London.

33 Mysterud, A. 1999. Seasonal migration pattern and home range of roe deer (Capreolus capreolus) in an altitudinal gradient in southern Norway. Journal of Zoology 247: 479–486.

34 Zimmermann, F. 2004. Conservation of the Eurasian Lynx (Lynx lynx) in a fragmented landscape - habitat models, dispersal and potential distribution. [s.n.], Lausanne. 179 p. pages.

35 Bates, D., M. Maechler, B. Bolker, S. Walker, R. H. B. Christensen, H. Singmann & B. Dai. 2014. lme4: Linear mixed-effects models using Eigen and S4. R package.

36 Sakamoto, Y., M. Ishiguro & G. Kitagawa. 1986. Akaike Information Criterion Statistics. Springer, Netherlands. pages.

37 Team, R. C. 2014. R: A language and Environment for statistical computing. Version 3.3.4 Very Secure Dishes. R Foundation for Statistical Computing, Vienna, Austria. https://www.r-project.org/.

38 Bartón, K. 2014. MuMIn: Multi-model interference. Version 1.15.6. R package, https://cran.r-project.org/web/packages/MuMIn/index.html.

39 Fiske, I., R. Chandler, D. Miller, A. Royle, M. Kery, J. Hostetler & R. Hutchinson. 2017. unmarked: Models for Data from Unmarked Animals. Version 0.11-0. R package, https://cran.r-project.org/web/packages/unmarked/index.html.

40 Roder, S. 2017. Red deer density drives wolf establishment in the Western Swiss Alps. MSc Thesis. Universitít Bern. 48 pages.

41 Akaike, H. 1974. A new look at the statistical model identification. IEEE Transactions on Automatic Control 19: 716–723.

42 Wilcoxon, F. 1945. INDIVIDUAL COMPARISONS BY RANKING METHODS. Biometrics Bulletin 1: 80–83.

43 Zengaffinen, N. 2016. Zweifel an Uni-Studie. Walliser Bote. December 1st, 2016.

44 Filliez, X. 2013. Les lynx et les loups, on le bousille, un point c’est tout. L’illustré 13: 20–23.

45 Valais, M. p. d. C. d. 2015. Ordonnance pénale du 12 novembre 2015 contre M. Lini Paccolat. MPB 15 1387.

46 Lüchtrath, A. & U. Schraml. 2015. The missing lynx - understanding hunters’ opposition to large carnivores. Wildlife Biology 21: 110–119.

47 Breitenmoser, U. 1998. Large predators in the Alps: the fall and rise of man’s competitors. Biological Conservation 83: 279–289.

48 Ceza, B., R. Kessler, K. Marti, N. Rochat & U. Tester. 2001. Wer tötet den Luchs?. 25, Pro Natura, Basel.

49 Moser, A. 2016. Rehbock, ledig, sucht…;. Netznatur. Swiss German Television (SRG1), Switzerland.

50 Norway, S. C. o. 2018. Motarbeidelse av rettsvesenet.. HR-2018-1784-A.

51 Norway, S. C. o. 2015. Straffutmåling for å ha hjulpet til med å skjule en ulovlig felt ulv. HR-2015-01134-A.

52 Project, W. J. 2019. Rule of Law Index. 978-0-9964094-1-4

53 Houte, A.-D. 2011. Les mutations de gendarmes depuis le XIX. siècle, entre contrainte institutionnelle et liberté individuelle. Travail et emploi: 29–39.

54 Schrami, U. 2019. Wildtiermanagement für Menschen. Pages 113–148 in H. M., editor. Wolf, Luchs und Bír in der Kulturlandschaft. Konflikte, Chancen, Lösungen im Umgang mit grossen Beutegreifern Eugen Ulmer KG, Stuttgart.

55 Gallant, J. C. & J. P. Wilson. 2000. Terrain Analysis: Principles and Applications in Primary topographic attributes. pages.

